# NODDI-derived measures of microstructural integrity in medial temporal lobe white matter pathways are associated with Alzheimer’s disease pathology and cognitive outcomes

**DOI:** 10.1101/2023.10.11.561946

**Authors:** Dana M. Parker, Jenna N. Adams, Soyun Kim, Liv McMillan, Michael A. Yassa, the Alzheimer’s Disease Neuroimaging Initiative

**Affiliations:** Department of Neurobiology and Behavior and Center for the Neurobiology of Learning and Memory, University of California, Irvine

**Author notes:** Corresponding authors: Dana M Parker, Michael A. Yassa. Data used in preparation of this article were obtained from the Alzheimer’s Disease Neuroimaging Initiative (ADNI) database (adni.loni.usc.edu). As such, the investigators within the ADNI contributed to the design and implementation of ADNI and/or provided data but did not participate in analysis or writing of this report. A complete listing of ADNI investigators can be found at: http://adni.loni.usc.edu/wp-content/uploads/how_to_apply/ADNI_Acknowledgement_List.pdf.

**Keywords:** diffusion tensor imaging, Neurite Orientation Dispersion and Density Imaging, Alzheimer’s disease, white matter, amyloid-beta, tau, memory, aging

## Abstract

**INTRODUCTION:** Diffusion tensor imaging has been used to assess white matter (WM) changes in the early stages of Alzheimer’s disease (AD). However, the tensor model is necessarily limited by its assumptions. Neurite Orientation Dispersion and Density Imaging (NODDI) can offer insights into microstructural features of WM change. We assessed whether NODDI more sensitively detects AD-related changes in medial temporal lobe WM than traditional tensor metrics.

**METHODS:** Standard diffusion and NODDI metrics were calculated for medial temporal WM tracts from 199 older adults drawn from ADNI3 who also received PET to measure pathology and neuropsychological testing.

**RESULTS:** NODDI measures in medial temporal tracts were more strongly correlated to cognitive performance and pathology than standard measures. The combination of NODDI and standard metrics exhibited the strongest prediction of cognitive performance in random forest analyses.

**CONCLUSIONS:** NODDI metrics offer additional insights into contributions of WM degeneration to cognitive outcomes in the aging brain.

## 1. Introduction

Alzheimer’s disease (AD) poses a significant public health challenge, which is exacerbated by the aging population. Currently it is estimated that roughly 6.7 million Americans are currently suffering from clinical AD with the potential of that number growing to 14 million in 2060 [1]. Emerging evidence indicates that development of pathology and neurobiological alterations in brain circuits occur long before the manifestation of symptoms, possibly even decades in advance [2–4]. Consequently, there is a pressing need to establish neuroimaging biomarkers with the ability to detect early neurobiological changes that contribute to the onset of cognitive deficits in AD.

White Matter (WM) tracts – large bundles of axons – enable communication between gray matter regions. These tracts are intriguing biomarkers in AD due to their vulnerability to Alzheimer’s pathology and subsequent impact on cognitive decline. The integrity of WM within the medial temporal lobe (MTL) may have particular relevance to detecting the earliest stages of AD. The MTL, encompassing the hippocampus, amygdala, and parahippocampal regions, plays a vital role in spatial and episodic memory [5,6]. Notably, the MTL is recognized as one of the earliest sites of atrophy [7] and tau tangle pathology [8]. Previous research has demonstrated strong relationships between gray matter integrity of the MTL, such as hippocampal volume [9,10], and disease progression. Further, MTL regions have been shown to have declining WM microstructure associated with normal aging [11]. Markers of MTL WM integrity in the earliest stages of AD have been less studied [12,13], though they may provide more sensitive information on structural decline.

Diffusion magnetic resonance imaging (dMRI), a noninvasive neuroimaging method, holds great promise in assessing the integrity of WM pathways. By employing dMRI, subtle and early changes in the brain’s WM can be detected [11,14]. Diffusion Tensor Imaging (DTI) can be used to calculate scalar indices derived from eigenvalues which are employed to describe anisotropy [11]. The most frequently utilized DTI indices include mean diffusivity (MD), which reflects the overall mean squared displacement of water molecules, and fractional anisotropy (FA), a measure of the magnitude of molecule diffusion that can be equated to anisotropic (i.e. directional) diffusion [11,15]. Both FA and MD are widely used as proxies for observing microstructural tissue changes during brain aging [16]. However, this tensor model poses limitations as it fails to capture the complex architecture of WM characterized by crossing, bending, twisting, and kissing fibers [17].

An alternative approach in dMRI involves tensor-free modeling, which encompasses various methods such as Q-shell imaging, Diffusion Kurtosis Imaging, and freewater elimination modeling [17,18]. Neurite Orientation Dispersion and Density Imaging (NODDI), another emerging tensor-free method, has the distinct advantage of allowing the examination of multiple compartments through multi-shell imaging, thereby offering a closer representation of the brain’s actual architecture [19–21]. Specifically, NODDI incorporates three microstructural compartments: intra-cellular, extracellular, and cerebrospinal fluid (CSF) [19]. NODDI generates metrics such as Neurite Density Index (NDI), which represents the portion of the tissue that is composed of axons and dendrites, and Orientation Dispersion Index (ODI), which reflects the angle variation and configuration of the neurite structure [22]. Previous research has validated NODDI measures against histological markers, revealing robust associations with compartment-specific indicators of neurite integrity. For example, Colgan et al. (2016) reported that NODDI, in a mouse model of human tauopathy (rTg4510), demonstrated the capacity to establish correlations within brain regions affected by tau. Remarkably, NDI outperformed the conventional FA metric by being the sole metric capable of detecting a correlation with tau burden in the hippocampus [23].

The objective of this study is to investigate whether NDI and ODI metrics, obtained through NODDI models of dMRI, exhibit comparable or improved performance of detecting AD progression in MTL WM. Given that the early stages of AD are characterized by memory loss, we focused on examining memory outcomes and pathology as they relate to potential early changes in major WM pathways within the MTL, specifically the cingulum, fornix, and uncinate fasciculus. Our goal was to determine if NODDI metrics enhance sensitivity in detecting microstructural changes in the brain, which may occur years before symptom onset in AD.

## 2. Methods

### 2.1 Data Source

Data used in the preparation of this article were obtained from the Alzheimer’s Disease Neuroimaging Initiative (ADNI) database (adni.loni.usc.edu). The ADNI was launched in 2003 as a public-private partnership, led by Principal Investigator Michael W. Weiner, MD. The primary goal of ADNI has been to test whether serial MRI, positron emission tomography (PET), other biological markers, and clinical and neuropsychological assessment can be combined to measure the progression of mild cognitive impairment (MCI) and early AD. For up-to-date information, see www.adni-info.org.

### 2.2 Study Participants

A total of 230 participants from the ADNI3 dataset were included in the study. Following an initial quality control stage, which involved assessing participants’ completion of scans and ensuring protocol dimension alignment and the use of a Siemens MRI machine for scan harmonization across sites, the final number of participants included in the analyses was 199. Due to a low number of participants with a dementia diagnosis (N=12), individuals diagnosed with MCI were combined with participants diagnosed with dementia for a Cognitive Impaired (CI) group (N=78), which were compared against a Cognitively Normal (CN) group (N=121). **Table 1** provides a breakdown of participant demographics.

**Table 1:** Demographic Breakdown.

|  | **CN (N=121)** | **CI (N=78)** |
| --- | --- | --- |
| **Age (range)** | 71(51-90) | 78(49-90) |
| **Sex: Female (%)** | 79(65) | 36(46) |
| **Years of Education (std)** | 17(2.36) | 15.8(2.5) |
| **Race** |  |  |
| **White (%)** | 94(78) | 64(82) |
| **Black (%)** | 19(16) | 8(10) |
| **Asian (%)** | 6(5) | 2(3) |
| **More than one (%)** | 2(2) | 3(4) |
| **Ethnicity** |  |  |
| **Hispanic/Latino (%)** | 2(2) | 3(26) |
| **Unknown (%)** | 1(0.8) | 1(1) |
| **Not Hispanic/Latino (%)** | 118(98) | 74(95) |
| **CDR-SB (std)** | .03(2.36) | 2.18(2.15) |
| **ADNI-MEM (std)** | 1.12(0.69) | 0.17(0.75) |

**Table 3.** Model performance evaluation metrics.

| Classification | Features | AUC<br>[CI] | Sensitivity (%)<br>[CI] | Specificity (%)<br>[CI] |
| --- | --- | --- | --- | --- |
| ADNI-MEM | NODDI | 0.65<br>[0.649 to 0.651] | 62<br>[0.619 to 0.621] | 65<br>[0.619 to 0.621] |
|  | Tensor | 0.67<br>[0.669 to 0.671] | 59<br>[0.589 to 0.591] | 65<br>[0.649 to 0.651] |
|  | Combination | 0.76<br>[0.759 to 0.761] | 69<br>[0.689 to 0.691] | 71<br>[0.709 to 0.711] |
| CDR-SB | NODDI | 0.69<br>[0.689 to 0.691] | 60<br>[0.599 to 0.601] | 69<br>[0.689 to 0.691] |
|  | Tensor | 0.70<br>[0.699 to 0.701] | 68<br>[0.678 to 0.682] | 70<br>[0.699 to 0.701] |
|  | Combination | 0.74<br>[0.739 to 0.741] | 63<br>[0.629 to 0.631] | 70<br>[0.699 to 0.701] |

AUC = Area under the curve, CI = 95% Confidence Intervals, ADNI-MEM = Alzheimer's Disease Neuroimaging Initiative Memory composite score, CDR-SB = Clinical Dementia Rating Scale Sum of Boxes

### 2.3 Structural and Diffusion MRI Processing

To ensure the homogeneity of the scan data across different sites, only scans obtained using Siemens MRI machines were selected for analysis. This decision aimed to minimize potential variations that could arise from using different machine manufacturers and ensure a more consistent and standardized dataset for the study. All participants were scanned using the ADNI 3.0 Advanced MRI scanning protocol, the first ADNI protocol to include multi-shell scan sequences. Each participant underwent a magnetization prepared rapid acquisition gradient echo (MPRAGE) sequence with the following parameters: echo time (TE)=2.98 ms, repetition time (TR)=2300 ms, inversion time=900ms, flip angle=10°, field of view (FOV)=208×240×256 mm3, acquired resolution =1×1×1mm. MRI parameter specifics can be found at http://adni.loni.usc.edu/methods/documents/mri-protocols/. T1-weighted images were corrected using Freesurfer 7 pipeline, which corrected for head motion and intensity inhomogeneity then the removal of non-brain tissue. After structural scans were quality checked, corrected hippocampal volume was calculated using intracranial volume and raw hippocampal volume Freesurfer output.

Diffusion scans were acquired via the following parameters: 232x232x160 mm @ 2x2x2 mm; TE=71 TR=3300. Three separate shells were collected at the following b-values: 500, 1,000, and 2,000 s/mm^2^ for a total of 112 directions. For the tensor metrics (FA and MD) the 1,000 s/mm^2^ shell was chosen as it was the b-value that was used in the single shell diffusion scan in the non-advanced ADNI3 protocol. Scans were preprocessed using MRtrix 3 [24](www.mrtrix.org). Gibbs ringing artifact [25], eddy current [26] and motion correction [27], were all completed prior to manual quality control. Tensor metrics were extracted via MRtrix3, and maps were created for FA and MD (**Figure 1A**). The Microstructure Diffusion Toolbox (MDT) [28] was used to calculate NDI and ODI maps (**Figure 1A**).

**Figure 1:**
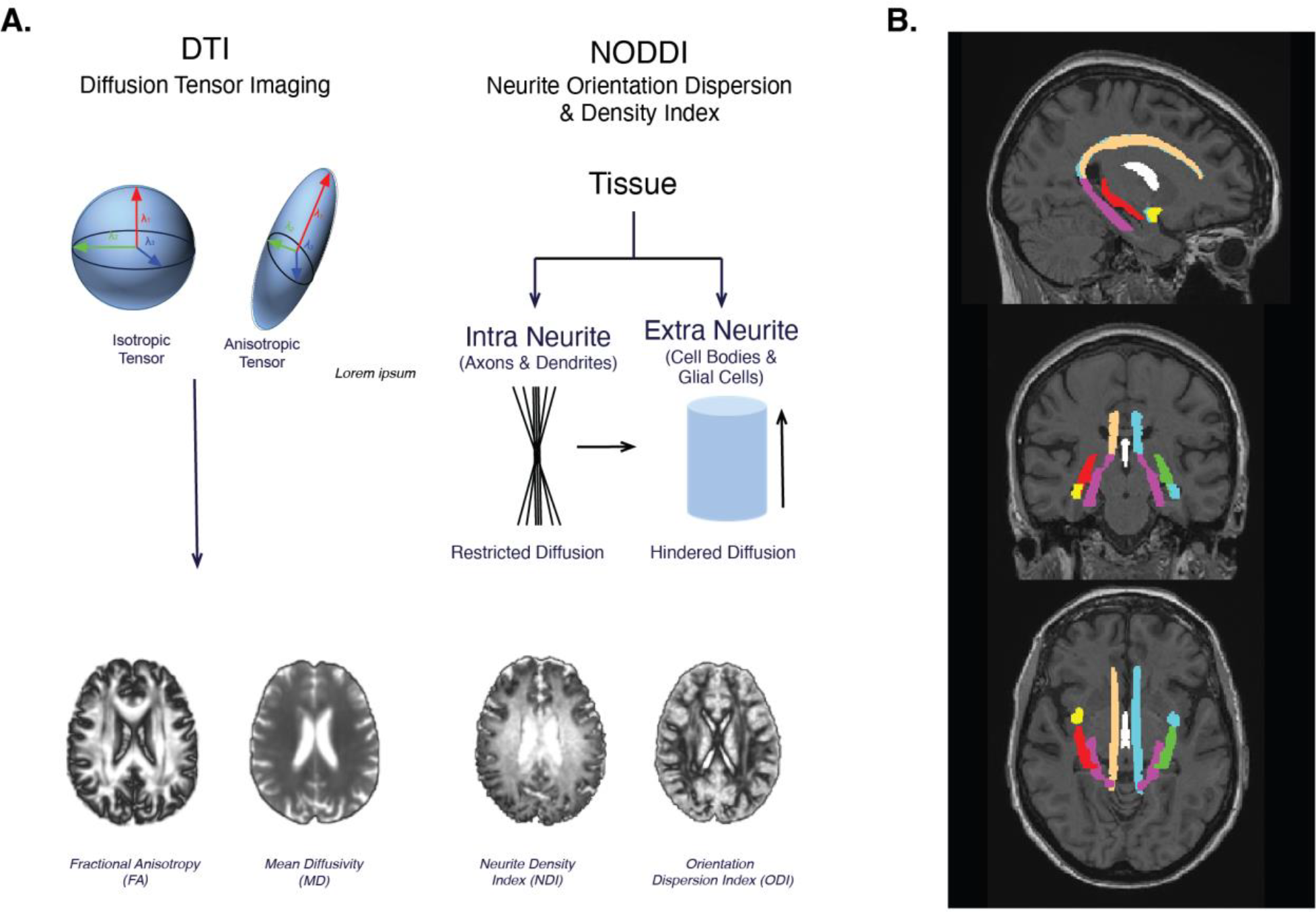
A. Overview of Diffusion Tensor imaging and overview of NODDI. B. Example of JHU WM Atlas ROIs

The following MTL ROIs from the JHU DTI based WM atlas (https://identifiers.org/neurovault.collection:264) were used in this analysis **(Figure 1B)**: cingulum cingulate, cingulum hippocampus, fornix column body, fornix ST and uncinate. Following preprocessing of the dMRI, images metrics for both NODDI and DTI were transformed to MNI space, and then the mean values of FA, MD, NDI, and ODI were extracted from each tract.

### 2.4 Clinical and Cognitive Measures

We analyzed the Clinical Dementia Rating Scale - Sum of Boxes score (CDR-SB), the Clinical Dementia Rating Scale - Memory (CDR-MEM) and ADNI-MEM as primary measures. The CDR-SB is a widely used clinical measure that assesses the severity of dementia in patients at a given time point. The “sum of boxes’’ component of the CDR provides enhanced precision for tracking changes over time and minimizes potential computational errors. The memory component of the CDR scale provides additional insights into the population based on their overall memory performance. Out of 199 included participants, 195 successfully completed the CDR and thus were included in the analyses of this measure [29].

To assess memory performance, we used the ADNI-MEM, a composite memory score derived from the cognitive battery as described by Crane et al. (2012) [30]. We opted to utilize this measurement as it provides a combined single memory score that serves as a robust and comprehensive representation of memory performance [30]. Out of the 199 participants, 186 had data available for ADNI-MEM.

### 2.5 Positron Emission Tomography

PET imaging protocols included 18F-Flortaucipir (FTP) to measure tau pathology and either 18F-Florbetapir (FBP) or 18F-Florbetaben (FBB) to measure Aβ pathology (depending on availability at time of collection). All preprocessing was completed by the ADNI PET core at UC Berkeley. Full details on PET processing and acquisition are available at the ADNI website https://adni.loni.usc.edu/wp-content/uploads/2012/10/ADNI3_PET-Tech-Manual_V2.0_20161206.pdf.

FTP data was analyzed between roughly 80-100 minutes post-injection across four 5-minute frames, using a Rousset approach for partial volume correction, and normalized using an inferior cerebellum gray reference region [31,32]. The mean standardized uptake value ratio (SUVR) values of FreeSurfer regions (version 7.1.1) were used. We focused on the mean SUVR of the entorhinal cortex and a meta-temporal ROI, which is a composite of the following regions: entorhinal, parahippocampal cortex, amygdala, fusiform, medial temporal, and inferior temporal [3].

FBP data was analyzed during a 50–70-minute post injection window across four 5-minute frames. A whole cerebellum reference region was used for normalization. Global Aβ was calculated via a cortical summary region which utilized Freesurfer-defined (version 7.1.1) frontal, anterior/posterior cingulate, lateral parietal, and lateral temporal regions [33,34]. To allow for the combination of the two tracers, FBP and FBB values were converted to centiloids [35].

### 2.6 Statistical Analysis

All statistical analyses were performed in Jamovi 2.3 (The Jamovi Project 2022) or Python (version 3.8). Correlation analyses performed were partial correlations controlling for age, sex, and education [36]. All p-values were corrected using the Bonferroni–Holm method in R (version 3.6.2) https://rdocumentation.org/packages/stats/versions/3.6.2 (correcting for 5 comparisons) using the p.adjust function from the “stats” package in R.

Random forest classification models were used to predict CDR-SB classification (CDR>0 vs CDR=0) and ADNI-MEM (mean split). Models were constructed as (1) tensor only, (2) NODDI only, and (3) combined models (best tensor, NODDI, and demographics). Random forest [37] is a type of ensemble machine learning algorithm that is widely used for classification or regression models including multi-dimensional variables [38]. We developed and implemented RF algorithms in Python using the Scikit-Learn library [39]. 100 decision trees consisting of different combinations of predictor variables were built for each model. Model performance was evaluated using area under the curve (AUC) of the receiver operating characteristic (ROC) curves, which was plotted from all the predicted values of the test data set using leave-one-out cross validation (LOOCV). The LOOCV design was specifically chosen given the small sample size [40] and built n classification models using n-1 subject each time which then utilized the classifier to determine the class of the left-out subject. Sensitivity and specificity were computed for each classification. We also performed bootstrapping (1000 simulations) to estimate 95% confidence intervals for the AUCs. Feature importance was also obtained using the Gini impurity index [37].

## 3. Results

### 3.1 NODDI measures show stronger associations with AD-related outcomes than standard tensor measures

We aimed to determine if using NODDI metrics could more sensitively account for complexities in WM tract microstructure, thus revealing stronger associations with outcome measures. Our two primary NODDI outcome measures were NDI, reflecting the density of how the neurites are packed, and ODI, reflecting orientation and directionality of the neurites (**Figure 1A**).

We first tested relationships between NODDI metrics of the MTL with clinical and cognitive performance, specifically the CDR-SB, CDR-MEM, and ADNI-MEM. Partial correlation analysis of NODDI results revealed that NDI consistently exhibited stronger correlations with outcome measures compared to ODI across all outcome measures, as shown in **Figure 2A** and **Supplementary Table 1**. Specifically, NDI in the cingulum cingulate and cingulum hippocampal regions demonstrated consistent and significant correlations with cognitive outcomes (ps_FDR_<0.05). In contrast, ODI only showed significant associations predominantly in the fornix and uncinate regions. Interestingly, ODI displayed consistent correlations across all three outcome measures (CDR-SB, ADNI-MEM, and hippocampal volume) in the fornix ST (ps_FDR_.001).

**Figure 2:**
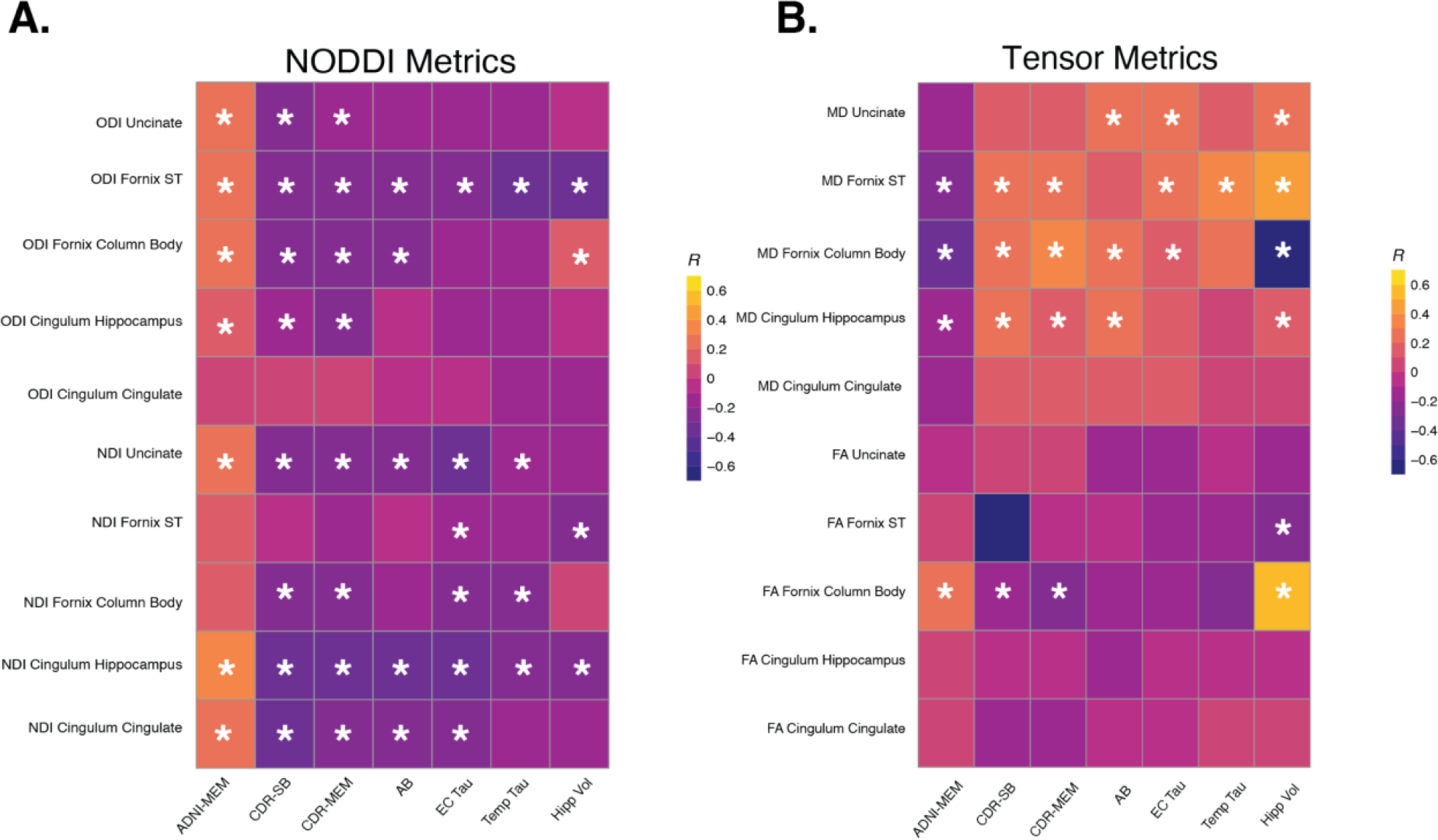
Correlation matrices between NODDI metrics (**A**) and tensor metrics (**B**) showing relationships between outcome measures and WM integrity of each ROI. NDI, neurite orientation index; ODI, * indicates p-value ≤ 0.05 after correction. Abbreviations: Fornix ST = Fornix Stria Terminalis, CDR-SB = Clinical Dementia Rating Scale Sum of Boxes, β-amyloid = Amyloid Beta, EC Tau = Entorhinal Tau, Temp Tau = Meta Temporal Tau, Hipp Vol = Hippocampal Volume.

We next tested associations between NODDI with Alzheimer’s pathology, specifically global Aβ, and both entorhinal and meta-temporal tau. Both NDI and ODI of various WM tracts exhibited significant correlations with pathology, with entorhinal tau demonstrating the greatest number of associations across both NODDI metrics. Entorhinal tau was strongly associated with NDI in the cingulum hippocampus (r=-0.375; p<0.001) and the uncinate (r=-0.373; p<0.001), and with ODI in the fornix ST (r =-0.288; p=0.004). Notably, ODI of the fornix ST showed a significant association in both tau ROIs (entorhinal r=-0.288; p<0.001; meta-temporal r=-0.319; p<0.001).

Finally, we investigated associations between NODDI measures and hippocampal volume as a proxy of neurodegenerative processes. Notably, hippocampal volume exhibited relatively fewer significant associations with the NODDI metrics. Hippocampal volume was only significantly associated with ODI of the fornix ST (r =-0.301; p<0.001) and NDI of the cingulum hippocampus (r =-0.226; p=0.0125) and fornix ST (r =-0.215; p=0.01).

We next examined the relationship between standard tensor-based diffusion metrics, namely FA and MD, with these same cognitive and pathology outcome measures (**Figure 2B**). Among the three cognitive outcome measures, there were no significant correlations with FA of any WM tracts assessed. On the other hand, MD of the cingulum hippocampus (r=-0.185; p=0.02), fornix ST (r=-0.273; p<0.01), and uncinate (r=0.244; p=0.01) demonstrated strong associations with ADNI-MEM. CDR-SB and CDR-MEM also showed similar results to ADNI-MEM (**Figure 2B**; **Supplementary Table 2A**). No other regions demonstrated significant findings in relation to MD.

We next assessed relationships between tensor metrics with AD pathology and hippocampal volume. There were no significant correlations between FA in any tract with either amyloid-beta or tau pathology in entorhinal cortex or meta-temporal ROI (ps_FDR_>0.075). However, FA of the fornix ST and fornix column body had significant relationships with hippocampal volume. MD across multiple tracts such as the fornix ST and fornix column body did show relationships with pathology and hippocampal volume, as indicated in **Figure 2B** and **Supplementary Table 2**.

Together, our results show that worse cognitive and pathological outcomes are associated with significant decreases in NDI and ODI, demonstrating the high sensitivity of NODDI metrics to detect these processes. In contrast, of the tensor metrics, increases in MD, but not FA, were associated with these measures, which highlights the reduced sensitivity of traditional DTI models.

### 3.2 Combination of NODDI and Tensor Metrics Best Predict Clinical/Cognitive Outcomes

Our next goal was to test which combination of diffusion metrics best predicted cognitive outcome measures associated with AD, and to determine the strength of prediction of these models, using ROC analyses. First, we constructed separate models to compare NODDI and tensor metrics directly. Second, we took the top metric from each model to create a combined diffusion model to maximize predictability. Due to the significant association between outcomes such as memory and demographic variables, final combined models included participants’ age, sex, and education.

We first tested prediction of memory performance using ADNI-MEM **(Figure 3A)**. Participants were divided into two using a median split method. Initial models were conducted separately for the NODDI metrics and tensor metrics, revealing distinct outcomes. The NODDI model (AUC=0.65; CI 95% bootstrap: [0.649,0.651]) displayed a more even distribution of Gini importance values across all included tracts and metrics (**Figure 3D**), whereas the tensor model (AUC=0.67 [0.669,0.671]) had a clear top performer, with a 29% performance gap between MD cingulum cingulate (71) and MD fornix column body (100), the two highest-ranking metrics on the importance table (**Figure 3C**). For the combined model, NDI cingulum hippocampus and MD fornix column body were selected along with the demographic variables. This combined model resulted in the strongest prediction, with an AUC of 0.76 [0.759,0.761].

**Figure 3:**
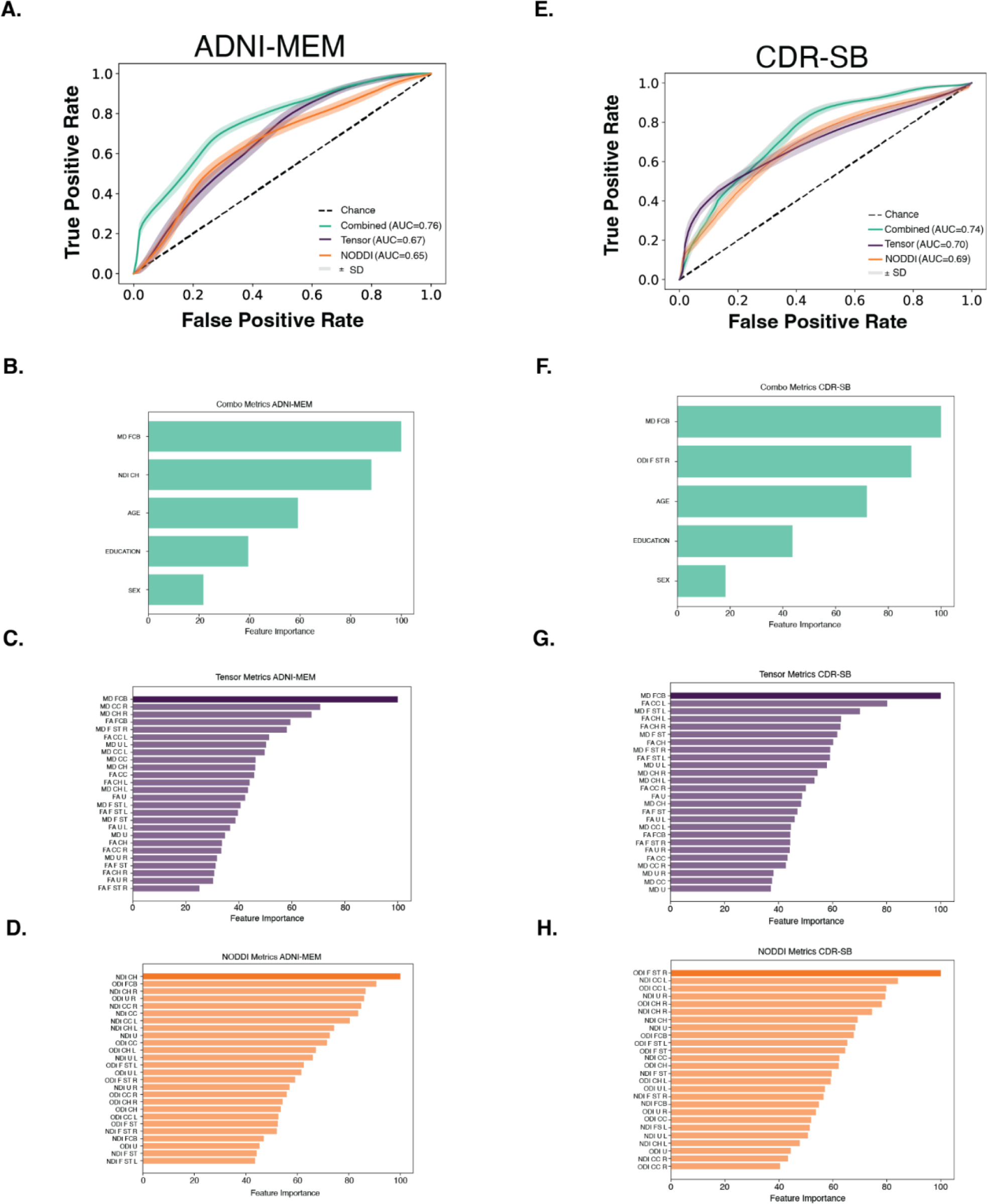
Characteristic curves for random forest classification models. A. ADNI-MEM model showing combined metrics (green), tensor metrics (purple) and NODDI metrics (orange). B. Gini Importance Table showing combined model results for ADNI-MEM. C. Gini Importance Table showing just tensor metric results for ADNI-MEM. D. Gini Importance Table showing only NODDI metrics results for ADNI-MEM. E. CDR-SB model showing combined metrics (green), tensor metrics (purple) and NODDI metrics (orange). F. Gini Importance Table showing combined model results for CDR-SB. G. Gini importance table showing only tensor metric results for CDR-SB. H. Gini Importance table showing only NODDI metrics results for CDR-SB. Dotted line = chance. Combination models included age, gender and education. Tensor and NODDI Gini Tables have top importance features bolded to show it was used in the combination modeling. Abbreviations: CC = Cingulum Cingulate, CH = Cingulum Hippocampus, FCB = Fornix Column Body, F ST = Fornix Stria Terminalis, U = Uncinate, R = Right hemisphere, L=left hemisphere.

We next tested prediction of the CDR-SB score (**Figure 3E**). We first classified participants as having a score indicating impairment (CDR-SB>0; n=80) or no impairment (CDR-SB=0; n=115). The NODDI model had an AUC of 0.69 [0.689,0.691]. Within this model, ODI of the fornix ST had the highest Gini value and was therefore the best predictor compared to the other NODDI tract metrics (**Figure 3H**). The tensor model had an AUC of 0.70 [0.699,0.701]. MD of the fornix column body had the highest Gini importance index (**Figure 3G**). The combined model, incorporating these top two performers, demonstrated an even higher predictive value for CDR-SB classification, yielding an AUC of 0.74 [0.739,0.741]. Specificity and sensitivity for each model can be found in **Supplementary Table 3**. CDR-MEM was not used as a model due to a lack of response diversity making it an insufficient outcome measure to be used with a machine learning model.

## 4. Discussion

The utilization of multishell diffusion imaging has opened new doors to understanding AD in ways that were previously unattainable through standard diffusion tensor modeling techniques. Previous studies using standard tensor models have left significant gaps in our understanding of the complexities of how WM microstructure is impacted by AD. Our research clearly highlights the advantages of employing the NODDI multishell compartmental model for gaining deeper insights into the pathology and cognitive outcomes of aging and AD. Our study demonstrates significant associations between NODDI and overall cognitive/clinical status, memory-related processes, as well as Alzheimer’s pathology. Overall, our study suggests that while DTI is a useful tool for studying aging, NODDI can give us additional information when it comes to cognition and pathology where tensor metrics such as FA fail to show significance.

Our study delved into the association between NODDI and tensor metrics with outcomes specific to aging and AD. The NODDI measure NDI, specifically when measured in the hippocampal cingulum, demonstrated more associations with cognition and pathology than tensor measures. In contrast, FA did not exhibit many associations with MTL WM integrity. NDI had strong correlative relationships specifically in the cingulum cingulate and cingulum hippocampal regions for tau and Aβ burden. Our results highlight significant associations between these NODDI metrics and tau pathology, with the NDI showing heightened sensitivity in the entorhinal region. Additionally, our results showed that Aβ had associations with the integrity of the cingulum, uncinate and fornix across both NDI and ODI metrics.

Although DTI has been widely employed to describe WM properties in previous research, it is not without limitations, such as issues related to crossing fibers [41]. It is widely recognized that as humans age, irrespective of the presence of neurodegenerative diseases, there is typically a reduction in fractional anisotropy (FA). However, FA can be a somewhat ambiguous metric, with the decline potentially attributable to various structural factors, including demyelination or a decrease in axon density [42]. While the development of alternative diffusion analysis pipelines has helped mitigate some of these limitations associated with DTI, the application of NODDI has emerged as a valuable method for assessing microstructural changes [41]. Our study demonstrates that FA alone exhibited weak, non-significant strength of association with nearly all the study’s outcome measurements. Recently, a study conducted on the ADNI3 cohort by Chen et al. (2023) [41] demonstrated that FA within specific MTL regions, particularly the hippocampal cingulum, exhibited correlations with tau burden. Our findings provide a more comprehensive examination of the compartmental complexity of WM, revealing that a stronger and more detailed association can be derived with NODDI than from the FA metric measurement alone. Based on this observation, we posit that NODDI metrics, with their capacity to discern alterations in microstructure, play a pivotal role in understanding the underlying changes occurring in WM during both the aging process and the progression of AD.

NDI may have increased sensitivity to detect early pathological changes compared to other metrics due to its ability to provide information at the cellular level. This information, which is achieved through NODDI’s use of multi-compartment analysis via the acquisition of multishell imaging, gives information at an anatomical level. This in turn is more nuanced and in depth when compared to the DTI methods which use a single diffusion shell to make more generalized statements about the status of water molecule movement within a region of tissue. Our study expands on previously published studies that investigated aging using NODDI and provides a more comprehensive assessment of how tensor metrics compare to multiple compartment approaches such as NODDI. One recent study found that both NODDI and tensor metrics are sensitive to age related changes in WM regions that are associated with aging [21]. While our work is similarly focused on the differences between these two diffusion analyses methods, our goal was to identify which measures were more associated with Alzheimer’s pathology (measured by PET) and related cognitive outcomes, and not the aging process per se. Another recent study investigated the relationship between flortaucipir PET and WM health focusing on NDI as their primary metric [25]. The authors showed that as Braak stages progressed, implying worsening levels of tau depositing, NDI in WM tracts decreased. Our findings build upon this previous work by demonstrating that utilizing a large multisite cohort study such as ADNI3 can allow for the extension of the previous work by comparing both cognitive and PET measurements within one study, with implications for follow-up studies to show longitudinality.

Finally, we tested the utility of NODDI and tensor metrics in predicting cognitive outcomes. We used the general rule that an AUC of 0.5 suggests no ability to predict, and 0.7 to 0.8 is acceptable prediction [43]. NODDI metrics, notably the NDI of the hippocampal cingulum, proved to be superior predictors of clinical outcomes when compared to demographic variables such as age, gender, and education. Similarly, tensor metrics, specifically the MD of the right fornix, exhibited acceptable predictive performance. The fusion of these metrics with demographic information yielded increased predictive models characterized with higher AUC values, suggesting models combining both NODDI and tensor metrics have increased sensitivity in the prediction of cognitive performance.

Currently, within the clinical trial and diagnostic realms, structural MRI serves as a crucial step in the process of confirming AD staging [3]. However, while volumetric measurements are employed to identify gray matter atrophy, the characterization of WM integrity can add more information about how neural communication and networks may be impacted. Our study underscores the value of integration of multishell diffusion acquisitions into the ADNI3 protocol, which has enabled a more complete understanding of the neurodegenerative processes occurring in both the aging population and individuals with AD. These metrics also open new avenues for future clinical trials, offering the potential to quantify early disease markers that were previously beyond reach using standard dMRI sequences. This advance holds promise for enhancing our ability to detect and address neurodegenerative conditions at earlier stages, potentially improving therapeutic interventions [3].

The study had several limitations that are important to note. First, the ADNI3 sample is not particularly diverse with respect to race and ethnicity. Our machine learning models would greatly benefit from a broader and more diverse dataset, as machine learning is not expected to generalize well to other samples if the training dataset is not sufficiently diverse [44]. Second, due to the sensitivity of neuroimaging to movement and imaging artifacts, our sample size was limited due to the need to censor participant data that did not pass quality control. Further, while our study focused on WM within the MTL, future investigations should use NODDI to assess WM integrity throughout the brain. Finally, neuroimaging studies investigating early stages of neurocognitive change with aging are limited by the relatively low sensitivity of neuropsychological and clinical measures of impairment. The use of more sensitive cognitive tasks is warranted in future studies.

In conclusion, our study offers new insights into the predictive potential of NODDI metrics in studies of AD. While tensor metrics have traditionally served as the standard for leveraging dMRI in assessing alterations in pathology and memory related to AD, our research suggests that conventional FA and MD measurements may miss more subtle microstructural changes. Notably, NDI emerges as an exceptionally sensitive metric, particularly in the context of entorhinal tau pathology and memory-related outcome measures. This underscores the potential of NODDI metrics to uncover subtle yet crucial insights into the complex dynamics of neurodegenerative processes.

## Acknowledgements

Data collection and sharing for this project was funded by the AD Neuroimaging Initiative (ADNI) (National Institutes of Health Grant U01 AG024904) and DOD ADNI (Department of Defense award number W81XWH-12-2-0012). ADNI is funded by the National Institute on Aging, the National Institute of Biomedical Imaging and Bioengineering, and through generous contributions from the following: AbbVie, Alzheimer’s Association; Alzheimer’s Drug Discovery Foundation; Araclon Biotech; BioClinica, Inc.; Biogen; Bristol-Myers Squibb Company; CereSpir, Inc.; Cogstate; Eisai Inc.; Elan Pharmaceuticals, Inc.; Eli Lilly and Company; EuroImmun; F. Hoffmann-La Roche Ltd and its affiliated company Genentech, Inc.; Fujirebio; GE Healthcare; IXICO Ltd.; Janssen Alzheimer Immunotherapy Research & Development, LLC.; Johnson & Johnson Pharmaceutical Research & Development LLC.; Lumosity; Lundbeck; Merck & Co., Inc.; Meso Scale Diagnostics, LLC.; NeuroRx Research; Neurotrack Technologies; Novartis Pharmaceuticals Corporation; Pfizer Inc.; Piramal Imaging; Servier; Takeda Pharmaceutical Company; and Transition Therapeutics. The Canadian Institutes of Health Research is providing funds to support ADNI clinical sites in Canada. Private sector contributions are facilitated by the Foundation for the National Institutes of Health (www.fnih.org). The grantee organization is the Northern California Institute for Research and Education, and the study is coordinated by the Alzheimer’s Therapeutic Research Institute at the University of Southern California. ADNI data are disseminated by the Laboratory for Neuro Imaging at the University of Southern California.

Special thanks to John Janecek for assistance with random forest computation and modeling. The authors are supported by NIA grants R01AG053555 (PI: M.A.Y), R21 AG075464 (PI: M.A.Y) and F32 AG074621 (PI: J.N.A).

## Supplemental Tables

**Supplementary Figure 1A:**
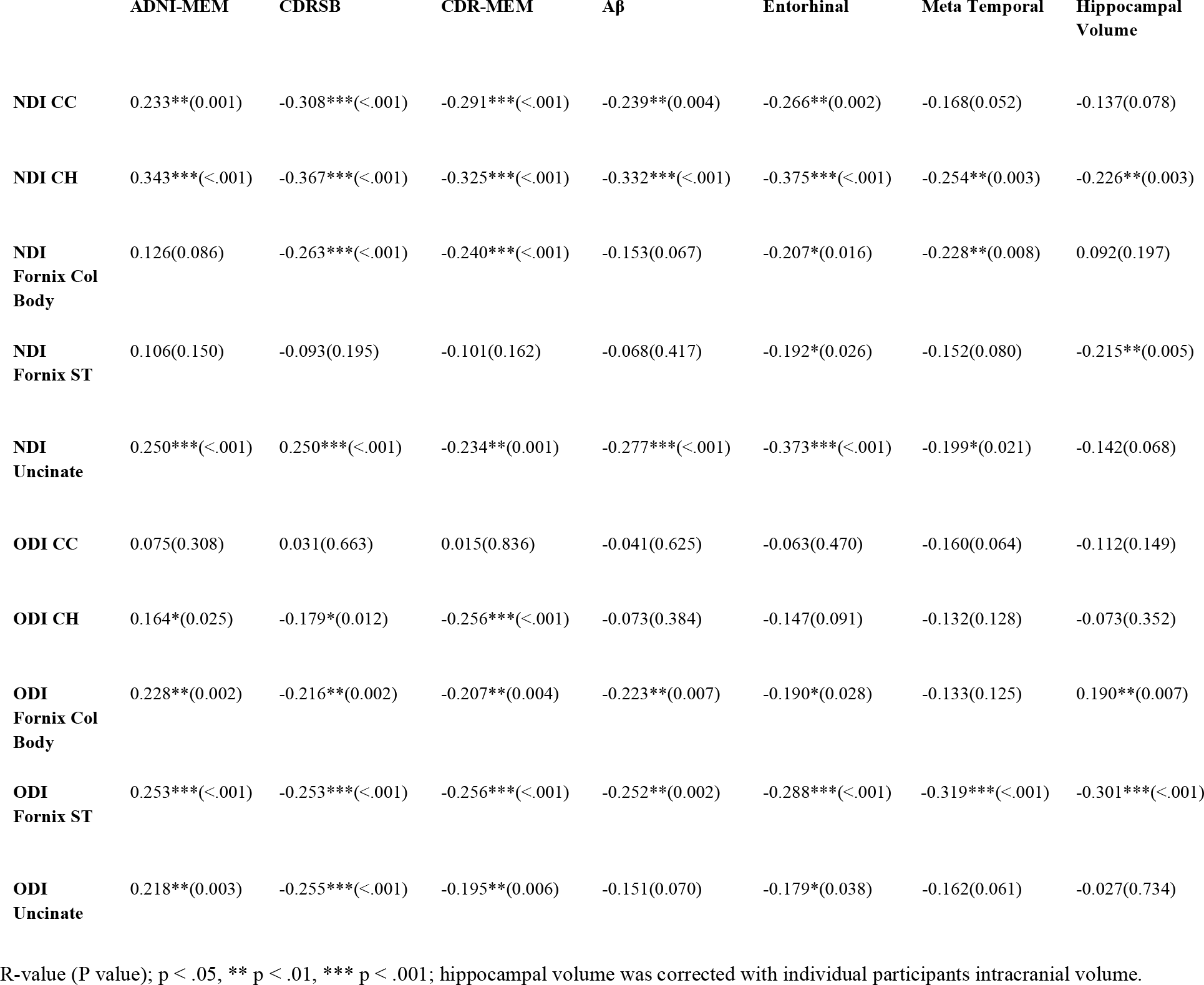
NODDI Metrics Partial Correlations with raw p-values.

**Supplementary Figure 1B:**
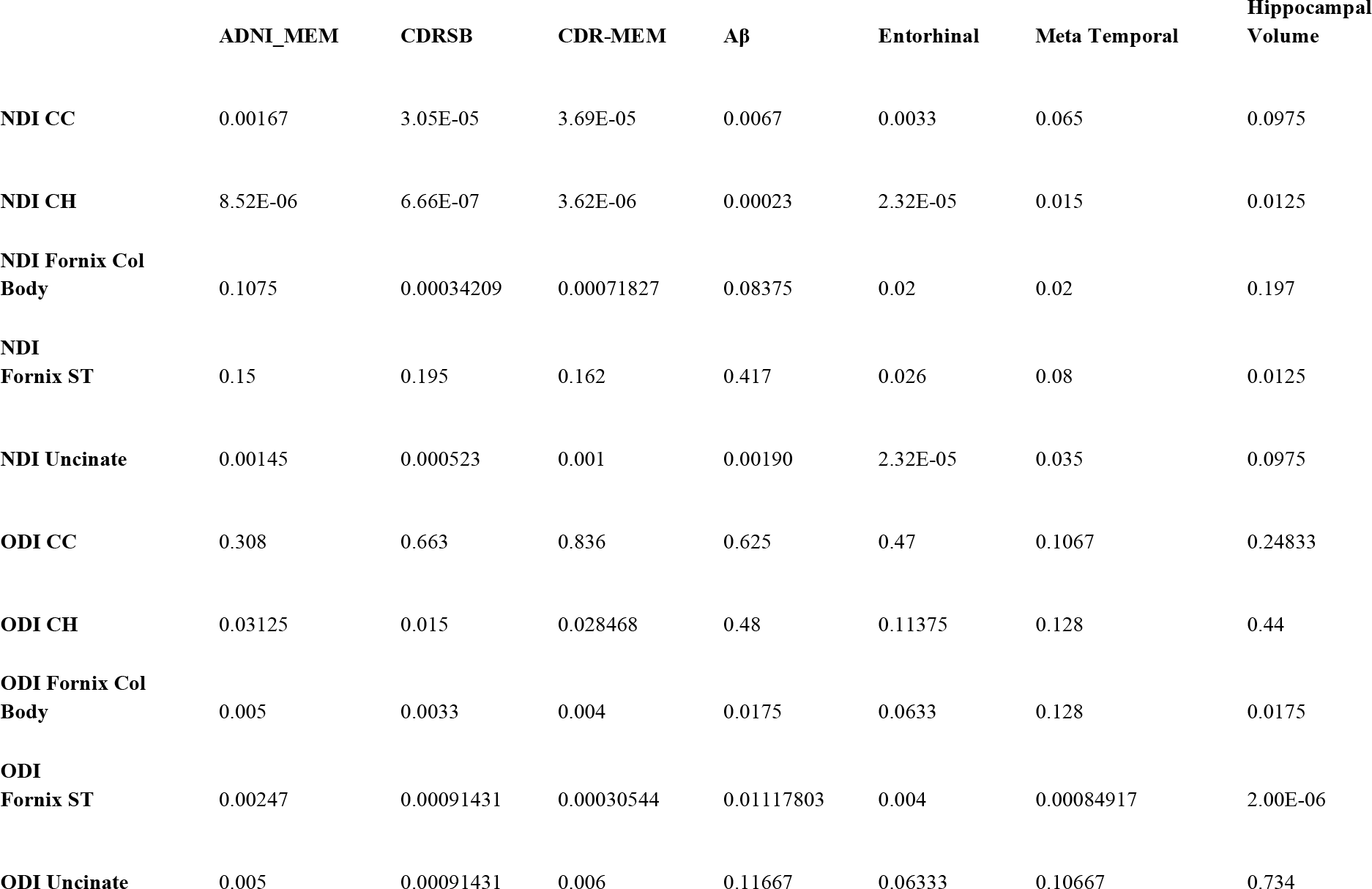
NODDI Corrected p-values.

**Supplementary Table 2A.**
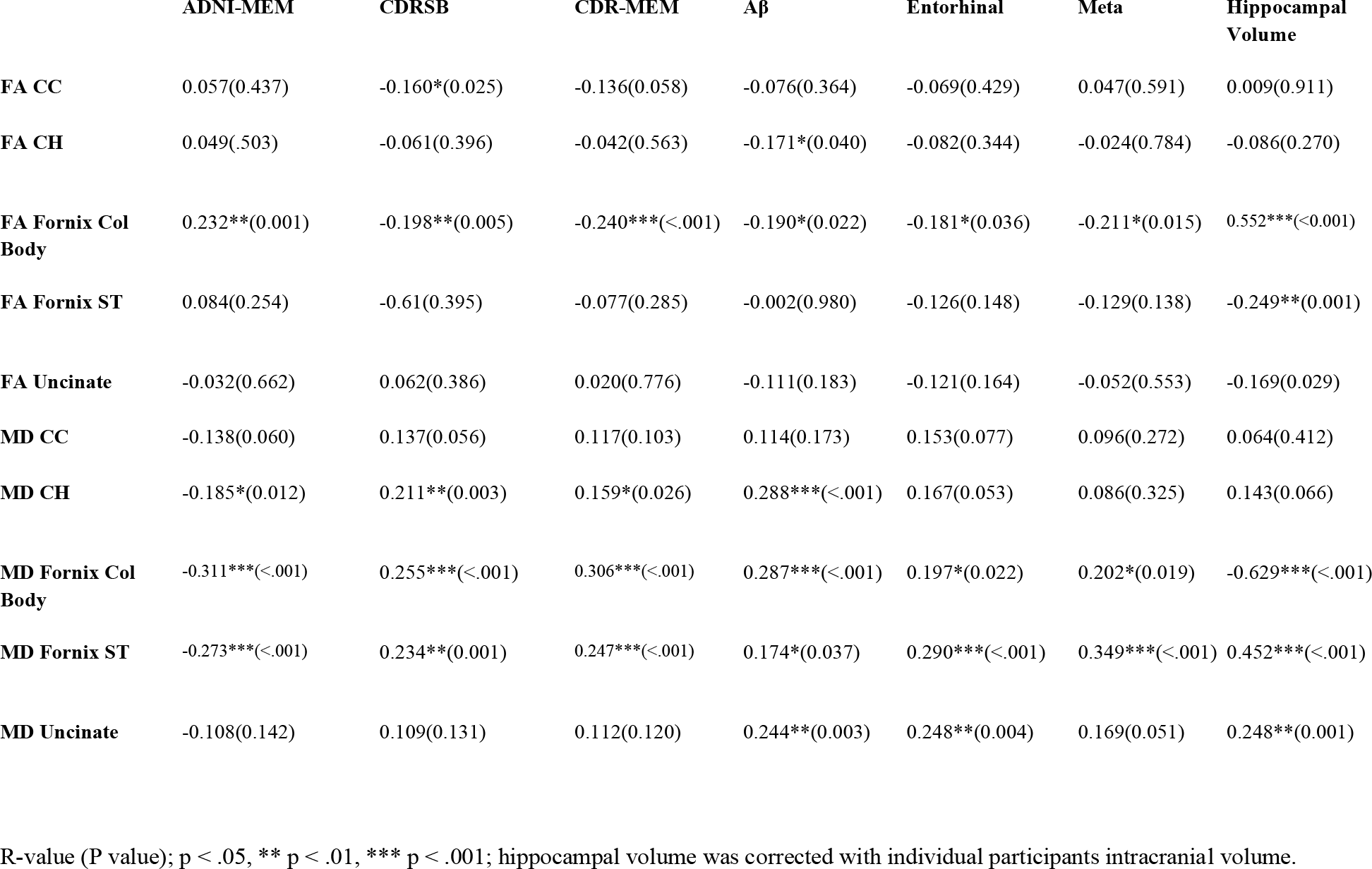
Tensor metrics partial correlations.

| | ADNI-MEM | CDRSB | CDR-MEM | Aβ | Entorhinal | Meta | Hippocampal Volume |
| --- | --- | --- | --- | --- | --- | --- | --- |
| FA CC | 0.057(0.437) | -0.160*(0.025) | -0.136(0.058) | -0.076(0.364) | -0.069(0.429) | 0.047(0.591) | 0.009(0.911) |
| FA CH | 0.049(.503) | -0.061(0.396) | -0.042(0.563) | -0.171*(0.040) | -0.082(0.344) | -0.024(0.784) | -0.086(0.270) |
| FA Fornix Col Body | 0.232**(0.001) | -0.198**(0.005) | -0.240***(<.001) | -0.190*(0.022) | -0.181*(0.036) | -0.211*(0.015) | 0.552***(<0.001) |
| FA Fornix ST | 0.084(0.254) | -0.61(0.395) | -0.077(0.285) | -0.002(0.980) | -0.126(0.148) | -0.129(0.138) | -0.249***(0.001) |
| FA Uncinate | -0.032(0.662) | 0.062(0.386) | 0.020(0.776) | -0.111(0.183) | -0.121(0.164) | -0.052(0.553) | -0.169(0.029) |
| MD CC | -0.138(0.060) | 0.137(0.056) | 0.117(0.103) | 0.114(0.173) | 0.153(0.077) | 0.096(0.272) | 0.064(0.412) |
| MD CH | -0.185*(0.012) | 0.211**(0.003) | 0.159*(0.026) | 0.288***(<.001) | 0.167(0.053) | 0.086(0.325) | 0.143(0.066) |
| MD Fornix Col Body | -0.311***(<.001) | 0.255***(<.001) | 0.306***(<.001) | 0.287***(<.001) | 0.197*(0.022) | 0.202*(0.019) | -0.629***(<.001) |
| MD Fornix ST | -0.273***(<.001) | 0.234**(0.001) | 0.247***(<.001) | 0.174*(0.037) | 0.290***(<.001) | 0.349***(<.001) | 0.452***(<.001) |
| MD Uncinate | -0.108(0.142) | 0.109(0.131) | 0.112(0.120) | 0.244***(0.003) | 0.248***(0.004) | 0.169(0.051) | 0.248***(0.001) |

R-value (P value); p < .05, \*\* p < .01, \*\*\* p < .001; hippocampal volume was corrected with individual participants intracranial volume.

**Supplementary Table 2B.** Tensor Metrics corrected p-values.

| | ADNI-MEM | CDRSB | CDR-MEM | Aβ | Entorhinal | Meta | Hippocampal Volume |
| --- | --- | --- | --- | --- | --- | --- | --- |
| FA CC | 0.62875 | 0.0625 | 0.058 | 0.455 | 0.429 | 0.73875 | 0.911 |
| FA CH | 0.62875 | 0.396 | 0.563 | 0.1 | 0.429 | 0.784 | 0.3375 |
| FA Fornix Col Body | 0.005 | 0.025 | 0.000729 | 0.1 | 0.18 | 0.075 | 1.43E-16 |
| FA Fornix ST | 0.62875 | 0.396 | 0.285 | 0.98 | 0.27333333 | 0.345 | 0.0025 |
| FA Uncinate | 0.662 | 0.396 | 0.776 | 0.305 | 0.27333333 | 0.73875 | 0.04833333 |
| MD CC | 0.075 | 0.07 | 0.103 | 0.173 | 0.077 | 0.325 | 0.412 |
| **MD CH** | 0.02 | 0.005 | 0.026 | 0.00116219 | 0.06625 | 0.325 | 0.0825 |
| **MD Fornix Col**<br>**Body** | 7.61E-05 | 0.0015797 | 1.35E-05 | 0.00116219 | 0.03666667 | 0.0475 | 1.34E-22 |
| **MD Fornix ST** | 0.00040567 | 0.0025 | 0.00050527 | 0.04625 | 0.00333226 | 0.00017659 | 9.44E-17 |
| **MD Uncinate** | 0.142 | 0.131 | 0.12 | 0.005 | 0.01 | 0.085 | 0.00166667 |

## Notes

**Conflict of Interest and Disclosure Statement:** M.A.Y. is Co-founder and Chief Scientific Officer of Augnition Labs, LLC. No other authors have conflicts of interest or disclosures.

### Competing Interest Statement

The authors have declared no competing interest.

